# Analysis of housekeeping genes in the peripheral blood of retinoblastoma patients

**DOI:** 10.1101/693101

**Authors:** Mayra Martínez-Sánchez, Jesús Hernandez-Monge, Mariana Moctezuma-Dávila, Martha Rangel-Charqueño, Vanesa Olivares-Illana

**Affiliations:** Laboratorio de Interacciones Biomoleculares y Cáncer. Instituto de Física, Universidad Autónoma de San Luis Potosí, Av. Manuel Nava 6. Zona Universitaria 78290 SLP, México; Cátedra CONACyT. Laboratorio de Interacciones Biomoleculares y Cáncer. Instituto de Física, Universidad Autónoma de San Luis Potosí, Av. Manuel Nava 6. Zona Universitaria 78290 SLP, México; División de Cirugía, Departamento de Oftalmología, Hospital Central “Ignacio Morones Prieto”, Universidad Autónoma de San Luis Potosí. SLP, México

**Keywords:** retinoblastoma, housekeeping gene, real time quantitative PCR, normalisation, internal control gene

## Abstract

Retinoblastoma is a pediatric neoplasia with a high incidence in non-developed countries. Nowadays the diagnosis is clinical and unfortunately in advanced stages of the diseases, which puts the child’s life risk. The search for molecular diagnosis on retinoblastoma is necessary. Due to their location and tendency to migrate, biopsies of retinoblastoma are not recommended then, peripheral blood samples can be a good source for the search of biomarker. RT qPCR is a sensitive method for gene expression quantification; in order to achieve optimal results, a crucial step is the reference gene choice that cannot be neglected under any circumstance. Six of the most commonly used housekeeping genes; GAPDH, HPRT1, B2M, TBP, RPL13a and 18S, were tested in the blood samples of patients diagnosed with retinoblastoma and healthy controls. The HPRT and TBP were found the most reliable genes whereas GAPDH, that is one of the most commonly used genes for normalisation, together with B2M, RPL13a and 18S have to be avoided. Using the selected reference genes, the *Rb* mRNA showed significant differences between patients and healthy children whereas no differences were found using to control groups. In the present study, we validate blood samples of patients with retinoblastoma.

## Introduction

Analysis of gene expression is widely used in studies with biological samples under different experimental conditions, or under disease conditions, and cancer is not an exception ^1^. A comparison of the expression levels of specific genes helps to have a more complete picture of certain diseases, to find the molecular markers of different kinds of cancers for prognosis and prediction, to detect DNA copy numbers, allelic discrimination, small non-coding RNA expression levels, and for virus detection, among other things ^2-7^. Real time quantitative polymerase chain reaction (qRT-PCR) is one of the most commonly used and powerful techniques for these types of studies ^5^. The results depend strongly on the normalisation of the data using an internal control gene with constant levels of expression ^8-10^. The internal control gene chosen, therefore, has to be stably expressed and show no difference in expression levels between groups, independent of the tissue type, cell line, experimental conditions, development stage, or disease state ^11-13^. However, it has been recently shown by many different groups that under certain non-physiological or experimental conditions, the levels of the most commonly used internal control or housekeeping genes in fact change, and are particularly affected in cancer ^12,14-20^. For example, studies validating housekeeping genes in glioblastoma show that TBP has been the most stable internal control whereas GAPDH shows significant differences between tumour samples and controls ^15,19^. In clear cell renal cell carcinoma, PPIA (peptidylprolyl isomerase A) and RPS13 (ribosomal protein S13) were the most stably expressed, whereas ACTB (β-actin), one of the most used housekeeping genes, was not a good choice for this carcinoma ^17^. Another example comes from a breast cancer housekeeping validation study, where PUM1 (Pumilio Homolog 1) was a better option as the internal control in this type of cancer, where B2M (β-2-microglobulin) and ACTB showed significant differences between tumour samples and controls ^18^. This demonstrates that even the most commonly used internal control genes have to be validated in order to select the appropriate housekeeping gene, depending on our experimental conditions.

Retinoblastoma is a common intraocular malignancy in the global paediatric population, being the second most common in children under four years of age. The incidence of the disease varies depending on the country or even the region. In developed countries the rates are between 2.2 to 6.2 per million habitants, while in non-developed countries, it can increase up to 24.5 per million habitants ^21,22^. Epidemiologic studies in the USA highlight a major incidence of retinoblastoma in the Latin community ^23^, nevertheless in Latin American countries the incidence rate of the disease is underestimated due to the lack of a national retinoblastoma registry and systematic statistics. Currently, the diagnosis of retinoblastoma is mainly clinical and in non-developed countries, where incidence is higher, it is performed in advanced stages of the disease, compromising the integrity of the eye and even the life of the patient ^21,24,25^. The most commonly used therapeutic modalities include chemotherapy, radiation therapy, cryotherapy, thermotherapy, enucleation (removal of the eyeball and optic nerve) and even orbital exetertion. The side effects of these treatments are important. Enucleation and exeteration for example, have major visual, cosmetic and psychological consequences due to the mutilation of the child. Without treatment, mortality from this tumour is 99%. It is clear that gaining information about the genetics is required in people affected by the disease in order to improve the diagnosis and treatment. Due to the nature and localisation of the tumour, a biopsy is not always an option for diagnosis of potential patients with the disease. With this in mind, we observe an urgent necessity to use non or less-invasive methods for diagnosis of this disease; hence blood samples might work as a good option for the development of molecular diagnostic techniques. To move forward on this topic, we examined the levels of some internal control genes for retinoblastoma in peripheral blood samples and compared them with healthy controls samples.

In the present work, we analysed the level of expression and stability of some of the most frequently used housekeeping genes in order to select and validate a proper internal control gene for peripheral blood samples of patients diagnosed with retinoblastoma. It has been reported the variation in the levels of those genes in tumour samples because of the intrinsic genetic instability of these cells. Here we show that in blood samples, four of six of the housekeeping genes we examined had shown very significant variation between the groups under study. The results presented here facilitates the utilization of blood samples in the retina cancer patients for RT-qPCR experiments for the diagnosis and prognosis of this disease.

## Results

### Expression Levels and stability analysis of potential housekeeping genes

We measured the expression levels and analysed the stability of potential housekeeping genes in order to find and validate an adequate internal control from the peripheral blood samples of paediatric patients with a diagnosis of retinoblastoma. We compared groups, patients and healthy controls, between 1 to 10 years old (**Table 1**). We choose six of the most commonly used housekeeping genes: β-2-microglobulin (B2M), Ribosomal protein L13a (RPL13A), Glyceraldehyde-3-phopsphate dehydrogenase (GAPDH), Hypoxanthine phosphoribosyltransferase 1 (HPRT1), 18S Ribosomal RNA (18S), and TATA-binding protein (TBP) (**Table 2**). The primers used are listed in Table 3, the efficiency of all primer sets was assayed by serial dilutions of cDNA and calculated according to the equation (1)^26^ (**Figure 1**). The amplicon sizes and specificity were checked on agarose gel electrophoresis (**Table 3 and Figure 2A**),

**Table 1.** Information of pediatric patients and controls.

| **Patient samples** | **Gender** | **Age (years)** | **Laterality** | **Age at diagnosis (months)** | **Control samples** | **Gender** | **Age (Years)** |
| --- | --- | --- | --- | --- | --- | --- | --- |
| **722828** | M | 9 | U | 6 | 1 | F | 1 |
| **899904** | M | 3 | B | 6 | 2 | F | 3 |
| **903921** | M | 3 | B | 2 | 3 | F | 7 |
| **920789** | M | 5 | U | 12 | 4 | M | 7 |
| **945305** | F | 2 | U | 9 | 5 | M | 7 |
| **945705** | M | 1 | U | 22 Days | 6 | M | 7 |
| **980172** | M | 2 | U | 18 | 7 | F | 9 months |
| **945705-2** | F | 2 | U | 19 | 8 | M | 2 |
|  |  |  |  |  | 9 | F | 10 |
|  |  |  |  |  | 10 | M | 10 |
|  |  |  |  |  | 11 | F | 10 months |

**Table 2.**
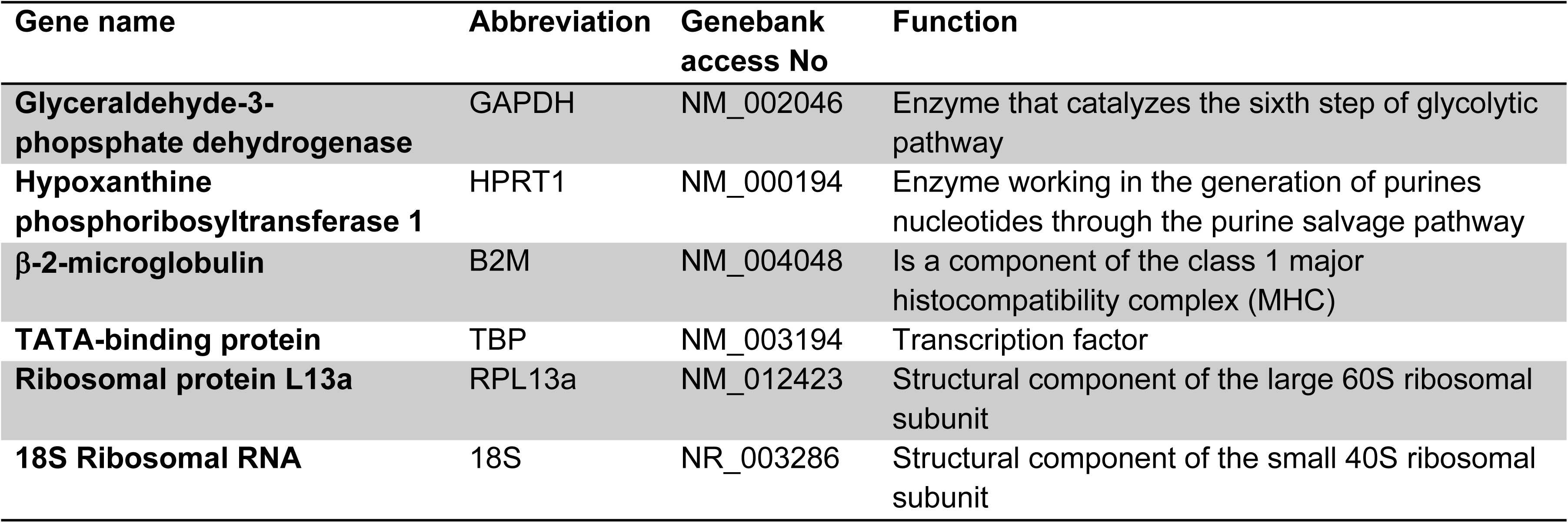
Housekeeping genes selected for expression analysis.

**Table 3.**
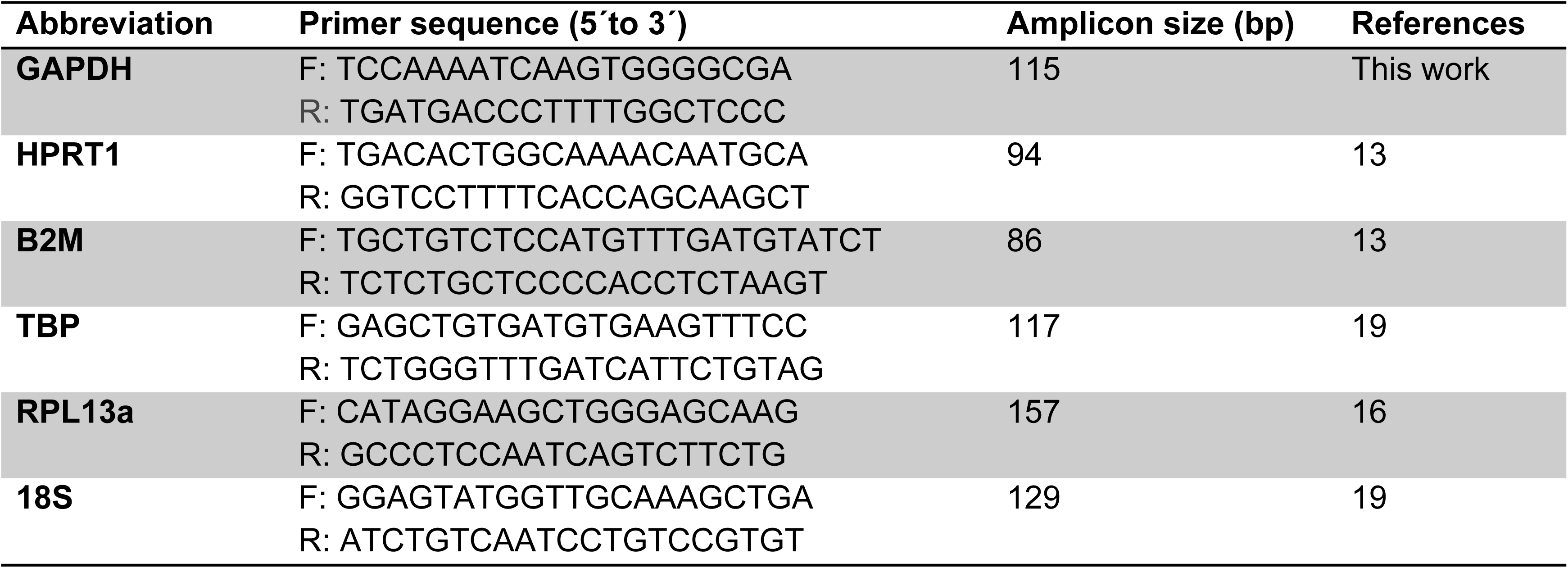
Primers sequence used for the real time pcr assay.

**Figure 1.**
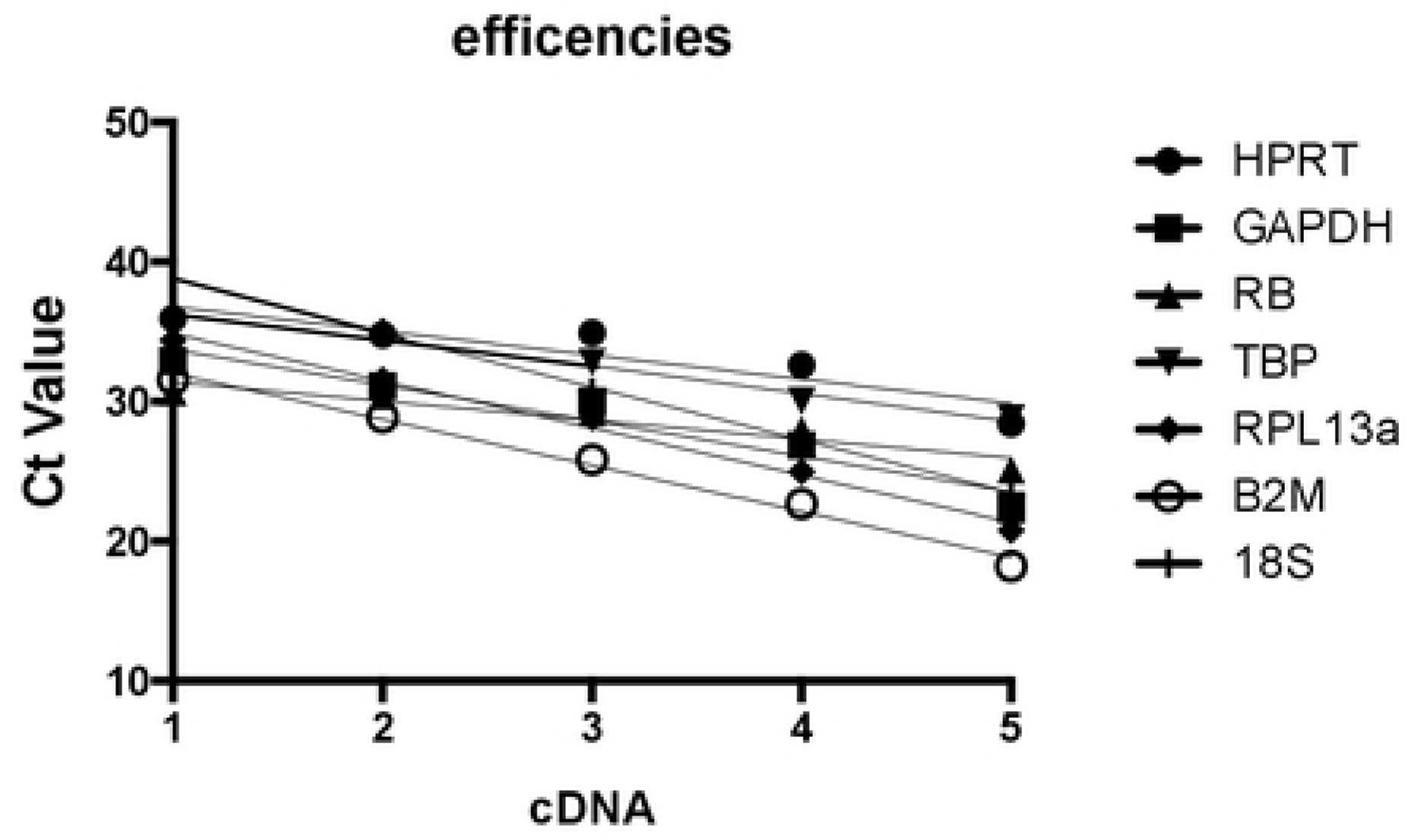
Determination of real-time PCR efficiencies of reference genes (HPRT, GAPDH, TBP, RPL13a, B2M, 18S) and target gene (Rb). The corresponding real-time PCR efficiencies were calculated according to equation (1) E=10^(−1/slope)^.

**Figure 2.**
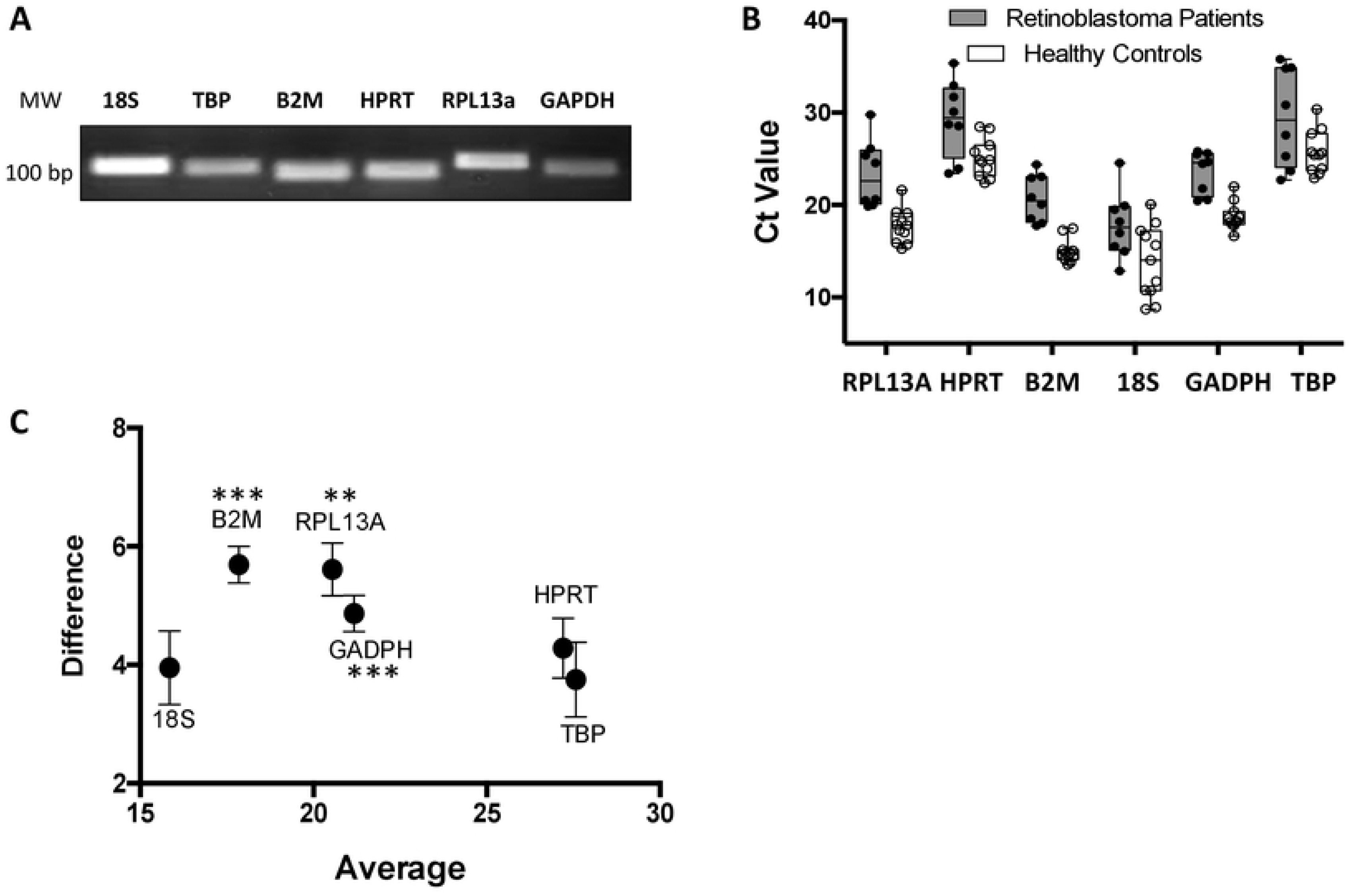
Comparison of expression levels of candidate housekeeping genes in blood samples of retinoblastoma patients and healthy controls. **(A)** Electrophoresis gel confirming the amplicon size and primer specificity used in the RT-qPCR for all the six housekeeping gene tested. **(B)** Boxes represent upper and lower quartiles of cycle threshold range with medians. Grey boxes correspond to retinoblastoma patients. White boxes represent healthy controls. **(C)** The differences of means and the matching symmetrical confidence intervals are shown for the relative expression of each housekeeping gene in the blood samples of patients and controls. If the symmetric confidence interval is included in the area of deviation and contains zero, the gene is considered equivalently expressed between patients and controls.

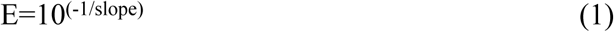

The transcriptional levels of the selected genes were determined in a panel of eight blood samples from patients with retinoblastoma and eleven blood samples of controls without a diagnosis of retinoblastoma. The six housekeeping genes analysed here showed a wide range of expression levels with Ct values between 9.8 to 35. The genes are distributed into three different categories: highly abundant genes such as 18S and B2M with Ct values between 9.8 and 24.5; genes with moderated abundance such as RPLI3A and GAPDH with Ct values between 15.2 and 29.7; and low abundance genes such as HPRT and TBP with Ct values between 22.7 and 35.1. The figure 2B shows raw Ct values of retinoblastoma patients (grey) and healthy controls (white). It could be observed that the amplitude in the expression levels is higher in patient samples than in the healthy controls. We found four to six cycles largest in the samples of patients with retinoblastoma compared with the samples of the controls, with the exception of GAPDH where the difference was just 0.4 cycles between the groups, suggesting a great variability in gene expression levels of blood samples of patients even in the so-called housekeeping gene (**Figure 2B**).

The results were first evaluated by using Mann-Whitney-Wilcoxon test. The analysis of the expression levels demonstrated that B2M, RPL13a and GAPDH have significant differences between the two groups suggesting that these genes have to be avoided as housekeeping in retinoblastoma blood sample patients. On the other hand, HPRT (0.0532), 18S (0.075) and TBP (0.267) showed no significant differences between samples (**Figure 2C and Table 4**). These commonly used statistical analysis shows that HPRT, TBP and 18S are the better option for an internal control gene for the blood samples of retinoblastoma patients in terms of expression levels.

**Table 4.** Statistical analysis of housekeeping level.

| Gene | Mann-Whitney-Wilcoxon | CV patients | CV controls | CV total |
| --- | --- | --- | --- | --- |
| **GAPDH** | 0.0014 ^ | 9.73 | 7.81 | 14.67 |
| **HPRT** | 0.0532 | 14.13 | 8.16 | 13.42 |
| **B2M** | 0.0003 ^ | 12.27 | 8.48 | 19,17 |
| **TBP** | 0.2670 | 18.14 | 8.88 | 14.47 |
| **RPL13a** | 0.0011 ^ | 15.64 | 10.31 | 18.85 |
| **18S** | 0.0750 | 20.28 | 28.21 | 26.29 |

Non-parametric Wilcoxon-Mann-Whitney test, highest values indicate best candidates ^ <0.05 indicates unsuitable housekeeping genes

The stability of the potential housekeeping genes was first examined by calculating the coefficient of variation (CV) of the Ct values for the retinoblastoma patient samples (CV patients), the healthy controls (CV controls) and for all of the biological samples (CV total). The expression levels of the housekeeping gene tested were less stable in patients (CV patients >10) blood samples compared with the controls (CV controls <10) except for 18S that shows more variation between the control samples (**Table 4**). The results showed HPRT, TBP and GAPDH were found to be the better-ranked genes. However, as we compared two experimental conditions (patients vs controls), the intergroup analysis (Mann-Whitney-Wilcoxon analysis table 4) is fundamental for the choice of the appropriated gene. Taking together, the results presented above revealed that GAPDH together with B2M and RPL13a present a significant difference between groups and were the worse ranked then, those were excluded for the further analysis.

Next, we tested 18S, TBP and HPRT genes using the online algorithm RefFinder ^27^. According to RefFinder, HPRT (1.00) was the best housekeeping gene to use, as the lowest geometric mean value represents the most stable and better-ranked gene, followed by TBP (1.8), and finally 18S (2.71) (**Table 5**). Using the geNorm software ^13^ we calculated the M value for these three genes, the M value is a reflect of the relative stability between all the reference genes analysed ^13^. GeNorm has a cut-off for an M value lower than 1.5, the better-ranked genes using this approach were HPRT and TBP (M=1.4) whereas 18S presented an M value much higher than the cut-off for geNorm (M=4.1) (**Table 5**). These results were corroborated using the NormFinder program, a mathematical model based on the variance estimation that correlated inversely to the expression stability; the lowest stability value indicates the more stable expressed candidate genes ^28^. NormFinder revealed that HPRT (0.016) followed by TBP (0.0249) are the more stable housekeeping genes whereas 18S (0.097) showed the highest value which meant that this was not recommended as good internal controls for normalisation. The best combination of two genes was HPRT and TBP (0.017) (**Table 5**).

**Table 5.**
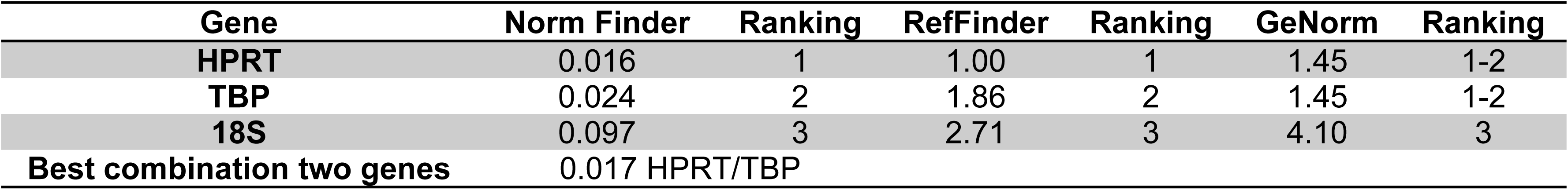
Stability ranking of the housekeeping genes.

Taken together, the expression levels and stability values, GAPDH, B2M, RPL13a, and 18S shouldn’t be used as internal control genes for blood samples of retinoblastoma patients. We could then conclude that HPRT and TBP are the better internal control genes for normalised gene expression data from blood samples of retina cancer patients, the use of the geometrical mean value of (TBP/HPRT) is also recommended to normalise the data.

### Validation of the identified housekeeping genes

To verify the pertinence of the selected housekeeping genes, normalised expression analysis was performed using expression data from *retinoblastoma* mRNA (*Rb* mRNA). In order to have two control groups to compare the levels of the *Rb* mRNA, we recruited 10 healthy adults control and analysed the six housekeeping genes in the new group. We compared the levels of the six reference genes between the two control groups (healthy adults vs children). The analysis revealed that the levels of these six housekeeping genes between the groups have significant differences between the control groups (**Supplementary Figure 1 and Supplementary Table 1**). These differences can only be attributed to the age, so far, only a couple of studies have compared the levels of proteins in blood samples between children and adults ^29,30^, in those cases, they observed also a significant differential expression levels of more than 100 proteins across the ages. Later we measure the levels of *Rb* mRNA into the three groups’ pediatric controls, patients and adults’ controls. A box plot showing the non-normalised expression levels for *Rb* mRNA was generated to show the general trends in expression levels of pediatric patients, pediatric controls and adult controls (**Figure 3A**). In supplementary figure 1 we observed that the levels of all the six housekeeping genes are a significant difference between controls pediatric and adults, however, we believe that the ratio between the *Rb* mRNA and the good reference gene has to remain constant between healthy children in comparison with healthy adults. Then we normalised the *Rb* mRNA expression levels in pediatric and adults controls groups with TBP, HPRT and the geometrical mean of the two better-ranked genes (TBP/HPRT) together with two of the most reference genes used in the literature as 18S and GAPDH. The data show that in fact, with the selected housekeeping genes the levels in the two control groups remain constant but the normalisation with 18S and GAPDH show a significant difference between the two control groups (**Figure 3B**).

**Figure 3.**
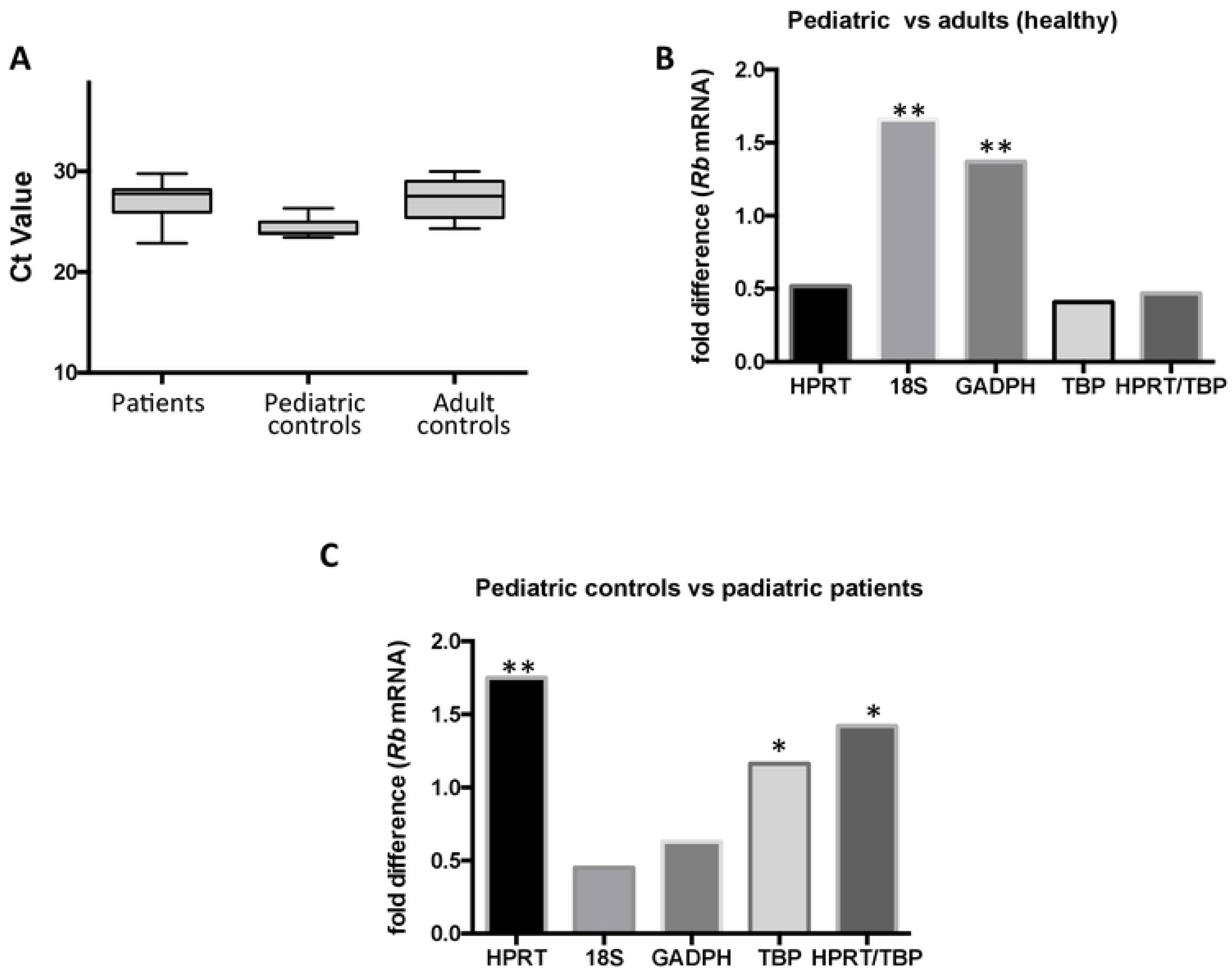
Effect of reference gene choice on Rt-qPCR normalization. **(A)** Box plots of averaged *Rb* mRNA expression data (Ct) for pediatric patients, controls and adult controls. Boxes represent upper and lower quartiles of cycle threshold range with medians. **(B)** The normalised relative expression ratio of *Rb* mRNA of pediatric and adults controls with TBP, HPRT, 18S and GAPDH. **(C)** The normalised relative expression ratio of *Rb* mRNA of pediatric patients and pediatric controls using as reference genes TBP, HPRT, 18S and GAPDH.

Finally, we normalized relative fold changes in expression of *Rb* mRNA between pediatric controls and pediatric patients using TBP, HPRT the geometrical mean of the two better-ranked genes (TBP/HPRT), together with 18S and GAPDH (**Figure 3C**). The normalization with HPRT, TBP and the TBP/HPRT showed significant differences in the expression levels of *Rb* mRNA between patients and controls; whereas GAPDH and 18S housekeeping genes were unable to detect these differences.

## Discussion

Housekeeping genes are ubiquitous and are involved in basic cellular functions, but it is evident from previous studies that the selection and validation of a housekeeping gene is a pre-requisite to the proper normalisation of gene expression data used in experimental conditions^14^, and is particularly important with genetically unstable cells, such as the tumour samples of cancer patients compared with healthy controls ^15,16,19^. It is not obvious, however, that the commonly used internal control genes are also altered in the blood samples of the patients with a very localised cancer such as retinoblastoma. Our results highlight that in this type of cancer, the expression of some genes are altered and has to be taken into consideration for all studies of gene expression analysis in blood samples.

Here we tested six of the most commonly used housekeeping genes: 18S, RPLI3A, B2M, HPRT, GAPDH and TBP. Our results show that TBP and HPRT are the better internal control genes for use in the blood samples of patients with retinoblastoma. We also suggested that the ideal normalization could be conducted by using the geometrical mean of TBP and HRT. Four of the six tested genes, 18S, B2M, GAPDH and RPL13a; were not optimal reference genes, and the use of some of them could generate erroneous results or can hide differences in gene expression.

The results presented here allow us to conclude that in retinoblastoma patients compared with healthy controls, even some of the so-called housekeeping genes are altered in peripheral blood. The *Rb* mRNA levels are also altered in retinoblastoma patients. These results provide evidence that the peripheral blood of retina cancer patients can be used for the search of potential biomarkers for the diagnosis and prognosis of this disease. Furthermore, we postulate that this alteration could also occur in other types of cancer where, as in retinoblastoma, the biopsy is not the best option, then blood samples can be potentially used. In conclusion, in blood samples of the paediatric population with retina cancer TBP, HPRT and the geometrical mean HPRT/TBP are better options to normalise the levels of gene expression of data generated from the qPCR analysis.

## Methods

### Samples and Ethics Statement

The study was developed within the framework of the approved protocol by the committee of research and ethics from the Central Hospital “Ignacio Morones Prieto” San Luis Potosí, SLP, Mexico. Diagnostic and therapeutic maneuvers were carried out according to the Official Mexican Standard NOM-012-SSA3-2012, which establishes the criteria for the execution of research projects for human health. The principles of the Helsinski Act of 1964 and its last revision in October 2013 are not transgressed. Written/signed informed consent was obtained from the guardians on behalf of the minors involved in the study. Ten peripheral blood samples were obtained from retinoblastoma patients and 21 healthy controls.

### Housekeeping Genes

Six of the most commonly used housekeeping genes were chosen for our study: Glyceraldehyde-3-phopsphate dehydrogenase (GAPDH), Hypoxanthine phosphoribosyltransferase 1 (HPRT1), Beta-2-microglobulin (B2M), TATA-binding protein (TBP), Ribosomal protein L13a (RPL13a) and 18S Ribosomal RNA (18S) (**Table 2**). The primers for HPRT1, RPL13a, 18S, TBP and B2M have been previously reported ^13,16,19^; GAPDH primers were designed previously in the laboratory (**Table 3**).

### RNA Extraction

250µl of mononuclear cells were isolated from peripheral blood and used to extract total RNA using the TRIzol method, according to the manufacturer’s protocol (Invitrogen). The precipitated RNA was resuspended in 50µl nuclease free water and treated with DNase. The concentration and A260/280 ratio of purified RNA were measured between 1.7 and 2.0 using a Nanodrop Spectrophotometer.

### cDNA Synthesis

The purified total RNA was reverse transcribed (1µg in each case) into cDNA in a reaction volume of 20 μl. The tubes were placed in the thermal cycler at 65°C for 5 minutes and then put on ice, and 8 µl of First Strand Mix (4 µl buffer first strand 5x, 1 µl ribolock, 1 µl M-MuLV Reverse Transcriptase enzyme and 2 µl dNTP’s [10mM]) were added. Tubes were placed in a thermal cycler at 4°C for 5 minutes, 60 minutes at 42°C, 72°C for 5 minutes and 4°C for 15 minutes. Three repetitions were performed.

### Electrophoresis analysis of the amplicons

The 6 housekeeping genes (18S, TBP, B2M, HPRT, RPL13a, GAPDH) were PCR amplified under the next conditions: denaturing temperature 95°C, melting temperature 60°C, extension temperature 68°C for 35 cycles, in a final volume of 25µl. The reactions were analysed through 1.5% agarose gel in Tris-EDTA buffer at 80V. The gel was stained in ethidium bromide and photodocumented.

### Quantitative Real-time PCR

RT-qPCR reaction was performed in 96-well microtiter plates. The amplification mixture consisted of 1 µl of each primer [10 µM], 20 ng of cDNA template and SYBR Green master mixed into a final volume of 12.5 µl. The PCR cycle conditions were set as follows: an initial denaturation step at 95°C for 10 minutes followed by 40 cycles of 95°C for 15 seconds, 60°C for 30 seconds, and 72°C for 30 seconds. Three replications were performed for each sample and each assay included a blank.

### Mathematical model for quantification in RT-qPCR

The mathematical model we used in order to quantified the target gene (Rb) in comparison with the housekeeping genes evaluated was proposed by Pfaffl M., in 2001^26^, and is based in equation (2). The efficiency corrected calculation model express the ratio of a target gen in samples vs controls in comparison to a reference gene.

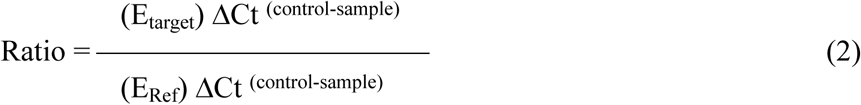

The E_target_ is the RT-qPCR efficiency of target gene transcript; E_Ref_ is the RT-qPCR efficiency of reference gene transcript. The efficiency was calculated according to the equation (1) (**Figure 1**).

### Data Analysis

The statistical analysis was performed using Graphpad Prism (version 6.0d GraphPad Software, CA). We selected the non-parametric Wilcoxon-Mann-Whitney for compared housekeeping genes levels. The expression stability of the potential housekeeping genes was evaluated by different methods. The coefficient of variation (CV) is used as a measure of expression stability. It was calculated by dividing the standard deviation of the Ct by the mean of the Ct of each group, the patients with diagnostic of retinoblastoma, the healthy controls and in all the samples. NormFinder is a method to identify stably expressed genes among a set of housekeeping genes. It is based on a mathematical model of gene expression that allows the estimation of the global variation of the candidate gene, and also the variation between the sample subgroups of a sample set ^28^. According to the NormFinder analysis, that best ranked is the one with the lowest stability value. RefFinder is a user-friendly web-based comprehensive tool that calculates a geometric mean based on the analysis of the most commonly used methods and programs currently available (Normfinder, BestKeeper, geNorm and the comparative ΔCt method) to assign a final ranking (http://leonxie.esy.es/RefFinder/ or http://150.216.56.64/referencegene.php?type=reference#) ^27^; geNorm ^13^. As in the case of NormFinder, the lowest value of the geometric mean is the top ranked.

## List of abbreviations

GAPDH: Glyceraldehyde-3-phopsphate dehydrogenase
HPRT1: Hypoxanthine phosphoribosyltransferase 1
B2M: Beta-2-microglobulin
TBP: TATA-binding protein
RPL13a: Ribosomal protein L13a
18S: 18S Ribosomal RNA
CV: coefficient of variation
ACTB: β-actin
RPS13: ribosomal protein S13
PUM1: Pumilio Homolog 1
PPIA: peptidylprolyl isomerase A
RTQ: Real time quantitative
PCR: polymerase chain reaction
SLP: San Luis Potosí

## Ethics approval

Research committee COFEPRIS 14 CI 24 028 083 and Ethics committee CONBIOETICA-24-CEI-001-20160427.

## Consent for publication

Not applicable

## Authors’ contributions

MM-S, MM-D and JH-M performed the experimental procedure of RNA extraction, cDNA synthesis, RT qPCR. MR-C obtained the blood samples of patients and controls and contributes to analyse the data. VO-I analyse the data and write the manuscript. All authors read and approved the final manuscript.

## Acknowledgments

We thanks to Yolanda Rebolloso-Gómez for technical support.

